# Metric Selection Effects in Consciousness Measurement: Lessons from Sleep EEG Analysis

**DOI:** 10.1101/2025.11.10.687628

**Authors:** Emma Dobbin

**Affiliations:** Consciousness Gradient Theory Group Ltd, United Kingdom

**Keywords:** consciousness measurement, sleep EEG, metric selection bias, circular reasoning, Lempel-Ziv complexity, spectral entropy, methodological validation

## Abstract

**Background:** Integrated Information Theory (IIT) predicts gradual consciousness changes, while Global Workspace Theory (GWT) emphasizes discrete threshold events. We tested whether sleep stage transitions exhibit these distinct patterns using quantitative EEG analysis.

**Methods:** We analyzed 622 sleep stage transitions from 12 healthy adults (Wake→N1, N1→N2, N2→N3, REM→Wake) using the Consciousness Gradient Index (CGI = √(φ × ρ) × 10). We employed two approaches: (1) an adaptive method selecting transition-specific metrics (alpha power, spindle density, spectral entropy), and (2) control analyses applying uniform metrics (spectral entropy, Lempel-Ziv complexity) across all transitions.

**Results:** The adaptive approach showed N1→N2 as the steepest transition (slope = −2.161, d = 2.814, p < 0.001). However, control analyses revealed this pattern to be metric-dependent. With spectral entropy applied uniformly, all transitions showed near-zero slopes (0.001-0.004), with N1→N2 ranking 3rd in magnitude. With Lempel-Ziv complexity, N1→N2 showed the smallest magnitude (0.001). Neither control supported N1→N2 as uniquely steep.

**Conclusions:** The apparent “dual architecture” resulted from selecting metrics based on known neurophysiological changes at each transition, creating circular reasoning. Control analyses using unbiased metrics showed no meaningful consciousness changes during any transition. This study demonstrates the importance of validation with uniformly-applied metrics and serves as a methodological warning about metric selection effects in consciousness research. The adaptive CGI framework successfully detected known neurophysiological changes but did not reveal fundamental differences in consciousness transition mechanisms.

## Introduction

Understanding how consciousness changes during sleep remains a fundamental challenge in neuroscience. Two major theoretical frameworks offer competing predictions: Integrated Information Theory (IIT) predicts gradual consciousness changes through continuous information integration (Tononi et al., 2016), while Global Workspace Theory (GWT) proposes discrete threshold mechanisms via broadcasting events (Dehaene & Changeux, 2011).

Sleep provides an ideal laboratory for testing these predictions, as individuals cycle through distinct states with well-characterized neural signatures. However, measuring consciousness quantitatively during these transitions has proven difficult. Traditional approaches rely on subjective reports or binary classifications (awake vs. asleep), failing to capture continuous gradients. More recent quantitative methods using complexity metrics (Casali et al., 2013) or connectivity measures (Tagliazucchi et al., 2016) have not systematically evaluated whether different transitions follow distinct patterns.

We developed the Consciousness Gradient Index (CGI = √(φ × ρ) × 10), combining information integration (φ) with adaptive response capacity (ρ). Our initial approach used an “adaptive” methodology, selecting different neural metrics for φ calculation based on transition-specific mechanisms: alpha power for arousal transitions, inverted spindle density for N1→N2, and spectral entropy for deep sleep transitions.

Here we report results from 622 sleep transitions showing that while this adaptive approach revealed strong patterns, control analyses using uniformly-applied metrics failed to replicate the findings. This demonstrates an important methodological trap in consciousness research: selecting metrics based on expected changes can create spurious patterns through circular reasoning.

## Methods

### Participants and Data

We analyzed polysomnographic recordings from 12 healthy adults (6 male, 6 female, age range: 25-35 years, mean age: 29.3 ± 3.2 years) from the Sleep-EDF Database Expanded (Kemp et al., 2000). All recordings were obtained from participants with no reported sleep disorders, neurological conditions, or psychoactive medication use. The database is publicly available and contains whole-night recordings at 100 Hz sampling frequency.

### EEG Recording and Preprocessing

Two EEG channels (Fpz-Cz and Pz-Oz) were used for all analyses to minimize contamination from respiratory or muscular artifacts. Raw EEG signals were bandpass filtered (0.5-30 Hz) using a zero-phase Butterworth filter. Sleep stages were scored in 30-second epochs according to Rechtschaffen and Kales criteria (Rechtschaffen & Kales, 1968). Hypnograms provided with the database were used to identify state transitions.

### Transition Detection and Analysis

We focused on four transition types: Wake→N1 (sleep onset), N1→N2 (spindle emergence), N2→N3 (deep sleep entry), and REM→Wake (awakening). For each transition, we extracted EEG data from 150 seconds before to 150 seconds after the transition point (total window: 300 seconds). CGI values were calculated in 30-second sliding epochs with 15-second overlap, yielding 19 measurements per transition. Linear slopes were fit using ordinary least squares regression to quantify the rate of change.

### Adaptive CGI Calculation

The formula CGI = √(φ × ρ) × 10 was applied with ρ = 1.0. The adaptive approach calculated φ differently for each transition:

- **Wake**→**N1 and REM**→**Wake:** φ from alpha power (8-13 Hz) using Welch’s method
- **N1**→**N2:** φ from inverted spindle density (11-16 Hz), with spindles detected via RMS power exceeding mean + 2 SD
- **N2**→**N3:** φ from spectral entropy (0.5-30 Hz)

All φ values were normalized to a 0-10 scale before CGI calculation.

### Control Analyses

To validate whether the adaptive pattern reflects genuine consciousness dynamics versus metric selection artifacts, we conducted two control analyses using established consciousness metrics applied uniformly:

### Control 1: Spectral Entropy (Uniform)

- Same spectral entropy calculation (0.5-30 Hz) applied to all four transition types
- No transition-specific modifications
- Shannon entropy of normalized power spectral density

### Control 2: Lempel-Ziv Complexity (Uniform)

- Binary sequence complexity analysis (Lempel & Ziv, 1976)
- EEG converted to binary (above/below median)
- Normalized complexity applied identically to all transitions

#### Validation Logic

If the dual architecture pattern is genuine, N1→N2 should remain the steepest transition with these unbiased metrics. If the pattern only appears with adaptive metrics, it indicates metric selection effects.

### Statistical Analysis

To address repeated measures structure, we employed subject-level analysis. For each subject and transition type, we calculated mean slopes across all detected transitions. Statistical comparisons used paired t-tests and repeated measures ANOVA on these subject-level means. Effect sizes were calculated using Cohen’s d. Significance was set at p < 0.05 (two-tailed).

## Results

### Dataset Characteristics

Across 12 subjects, we analyzed 622 transitions: 136 Wake→N1 (11.3 per subject), 207 N1→N2 (17.2 per subject), 247 N2→N3 (20.6 per subject), and 32 REM→Wake (4.6 per subject in 7 subjects).

### Adaptive Method Results

Subject-level analysis showed distinct patterns:

- Wake→N1: Mean slope = −0.706 ± 1.084
- N1→N2: Mean slope = −2.161 ± 2.859
- N2→N3: Mean slope = −0.503 ± 1.094
- REM→Wake: Mean slope = +0.333 ± 0.831

N1→N2 showed the steepest magnitude, approximately 3-fold greater than other NREM transitions.

#### Statistical Comparisons

Repeated measures ANOVA revealed highly significant differences (F = 45.564, p < 0.001).

Paired t-tests comparing N1→N2 against other transitions:

- N1→N2 vs. Wake→N1: t(11) = −9.334, p < 0.001, d = −2.814
- N1→N2 vs. N2→N3: t(11) = −6.341, p < 0.001, d = −1.912

These extremely large effect sizes suggested N1→N2 operates through a qualitatively different mechanism.

### Control Analysis Results

To validate whether this pattern reflects genuine consciousness dynamics, we applied spectral entropy and Lempel-Ziv complexity uniformly across all 622 transitions.

#### Spectral Entropy Control (Applied Uniformly)

- Wake→N1: Mean slope = 0.004 ± 0.009
- N1→N2: Mean slope = 0.002 ± 0.007
- N2→N3: Mean slope = −0.002 ± 0.005
- REM→Wake: Mean slope = −0.001 ± 0.008

#### Lempel-Ziv Complexity Control (Applied Uniformly)

- Wake→N1: Mean slope = 0.001 ± 0.006
- N1→N2: Mean slope = 0.001 ± 0.006
- N2→N3: Mean slope = −0.004 ± 0.010
- REM→Wake: Mean slope = −0.001 ± 0.003

Both controls showed slopes near zero (0.001-0.004), indicating no meaningful consciousness changes during any transition when using unbiased metrics. N1→N2 was not the steepest transition with either control method. In fact, with spectral entropy, N1→N2 ranked 3rd in magnitude, and with Lempel-Ziv, it showed the smallest magnitude.

## Discussion

### Summary of Findings

Our adaptive approach initially suggested a “dual architecture” where N1→N2 showed threshold-like behavior (steep slope, large effect size d = 2.814) while other transitions showed gradual changes. However, control analyses using uniformly-applied metrics contradicted this pattern. When spectral entropy and Lempel-Ziv complexity—two established consciousness proxies—were applied identically across all transitions, we observed essentially no consciousness changes (slopes ∼0.001-0.004) during any transition, and N1→N2 was not uniquely steep.

### The Circular Reasoning Trap

The control analyses revealed a critical methodological flaw in our adaptive approach. The apparent “dual architecture” resulted from selecting metrics **because** they change at specific transitions:

1. **Spindles naturally emerge during N1**→**N2** → We measured spindle density → Found steep changes → Claimed threshold mechanism
2. **Alpha naturally declines during wake**→**sleep** → We measured alpha power → Found gradual changes → Claimed gradual integration
3. **Complexity naturally decreases into deep sleep** → We measured spectral entropy → Found gradual changes → Claimed IIT validation

This is circular reasoning: selecting metrics based on known neurophysiological changes, then claiming to discover those changes represent consciousness dynamics. The adaptive method’s strong statistical results (d = 2.8) reflect well-documented EEG phenomena— spindle emergence, alpha decline, complexity reduction—not fundamental differences in how consciousness transitions operate.

### What We Actually Measured

The adaptive CGI framework successfully detected known neurophysiological changes:

- Spindle emergence during N1→N2 (well-established since the 1960s)
- Alpha power decline during sleep onset (well-established)
- Spectral entropy reduction into deep sleep (well-established)

However, these neurophysiological changes do not necessarily reflect consciousness-level changes. The control analyses demonstrate this distinction: when unbiased consciousness metrics were applied uniformly, they showed no meaningful transitions at all.

### Implications for Consciousness Measurement

Our findings highlight a subtle but pervasive problem in consciousness research: **the temptation to select metrics that “work”**. Even when theoretically motivated (as our adaptive approach was), metric selection based on expected patterns can create spurious findings.

The solution is straightforward but often neglected: **validation with uniformly-applied metrics**. Any claim that different conditions show different consciousness dynamics must be tested with at least one metric applied identically across all conditions. If the pattern only appears with condition-specific metrics, it likely reflects those specific neural features rather than consciousness per se.

### Reconciling IIT and GWT

Our initial goal was to reconcile IIT and GWT by showing both operate in different contexts. The control analyses suggest a different conclusion: the apparent conflict between these frameworks may partly reflect how researchers select metrics to test them. IIT studies often use complexity/integration measures; GWT studies often focus on broadcasting/threshold events. If both approaches inadvertently select metrics that confirm their predictions, apparent differences may be methodological rather than theoretical.

This does not invalidate either framework—both make important predictions about consciousness mechanisms. However, it suggests that testing these predictions requires more rigorous control of metric selection bias than has been typical in the field.

### Methodological Recommendations

Based on our findings, we recommend:

1. **Always include uniform-metric controls** when comparing consciousness across conditions
2. **Pre-register analysis plans** specifying metrics before data collection
3. **Report negative controls** when uniform metrics fail to replicate adaptive findings
4. **Distinguish neurophysiology from consciousness** - EEG changes don’t automatically equal consciousness changes
5. **Be transparent about metric selection rationale** and acknowledge when it’s post-hoc

### Limitations

Several limitations warrant consideration. First, our sample consisted of healthy young adults (age 25-35); generalizability requires testing in diverse populations. Second, we analyzed only four transition types; additional transitions (e.g., N2→REM, N3→N2) should be examined. Third, our control analyses used only two uniform metrics; testing additional unbiased measures would strengthen conclusions. Fourth, the REM→Wake analysis included only 32 transitions from 7 subjects, limiting statistical power for that comparison.

Most importantly, our negative controls do not prove consciousness doesn’t change during sleep transitions—only that our methods failed to detect it. Better metrics or methods may succeed where ours failed. However, the burden of proof now lies on demonstrating this with properly controlled validation.

### Future Directions

Future research should:

- Test additional uniform metrics (permutation entropy, global connectivity measures)
- Apply similar control analyses to other consciousness contexts (anesthesia, meditation)
- Develop metrics specifically validated against subjective reports across multiple conditions
- Create standardized benchmarks for consciousness measurement validation

The field would benefit from consensus criteria for what constitutes valid consciousness measurement, analogous to validation standards in other measurement domains.

## Conclusion

We explored whether sleep stage transitions exhibit a “dual architecture” reconciling IIT and GWT predictions. An adaptive approach selecting transition-specific metrics suggested such a pattern, with N1→N2 showing uniquely steep changes. However, control analyses using uniformly-applied consciousness metrics (spectral entropy, Lempel-Ziv complexity) failed to replicate this finding, showing essentially no consciousness changes during any transition.

This demonstrates that apparent patterns in consciousness measurement can result from metric selection effects rather than genuine differences in consciousness dynamics. The adaptive CGI framework successfully detected known neurophysiological changes but did not reveal fundamental differences in transition mechanisms. Our findings serve as a methodological warning: even theoretically-motivated metric selection requires validation with uniformly-applied controls to avoid circular reasoning.

The study contributes important methodological insights about validation requirements in consciousness research while highlighting the distinction between measuring neural activity changes and measuring consciousness itself.

